# Swimming, Feeding and Inversion of Multicellular Choanoflagellate Sheets

**DOI:** 10.1101/2023.03.14.532283

**Authors:** Lloyd Fung, Adam Konkol, Takuji Ishikawa, Ben T. Larson, Thibaut Brunet, Raymond E. Goldstein

**Affiliations:** Department of Applied Mathematics and Theoretical Physics, Centre for Mathematical Sciences, University of Cambridge, Wilberforce Road, Cambridge CB3 0WA, United Kingdom; Department of Biomedical Engineering, Tohoku University, 6-6-01 Aoba, Aramaki, Aoba-ku, Sendai 980-8579, Japan; Department of Biochemistry & Biophysics, University of California, San Francisco, 600 16th St., San Francisco, CA 94143-2200, USA; Department of Cell Biology and Infection, Institut Pasteur, 25-28 rue du Dr. Roux, 75724 Paris Cedex 15, France

## Abstract

The recent discovery of the striking sheet-like multicellular choanoflagellate species *Choanoeca flexa* that dynamically interconverts between two hemispherical forms of opposite orientation raises fundamental questions in cell and evolutionary biology, as choanoflagellates are the closest living relatives of animals. It similarly motivates questions in fluid and solid mechanics concerning the differential swimming speeds in the two states and the mechanism of curvature inversion triggered by changes in the geometry of microvilli emanating from each cell. Here we develop fluid dynamical and mechanical models to address these observations and show that they capture the main features of the swimming, feeding, and inversion of *C. flexa* colonies.

---

Some of the most fascinating processes in the developmental biology of complex multicellular organisms involve radical changes in geometry or topology. From the folding of tissues during gastrulation [1] to the formation of hollow spaces in plants [2], these processes generally involve coordinated cell shape changes, cellular division, migration and apoptosis, and formation of an extracellular matrix (ECM). It has become clear through multiple strands of research that evolutionary precedents for these processes exist in some of the simplest multicellular organisms such as green algae [3, 4] and choanoflagellates [5], the latter being the closest living relatives of animals. Named for their funnel-shaped collar of microvilli that facilitates filter feeding from the flows driven by their beating flagellum, choanoflagellates serve as model organisms for the study of the evolution of multicellularity.

While well-known multicellular choanoflagellates exist as linear chains or “rosettes” [6] held together by an ECM [7], a new species named *Choanoeca flexa* was recently discovered [8] with an unusual sheet-like geometry (Fig. 1) in which hundreds of cells adhere to each other by the tips of their microvilli, without an ECM [9]. The sheets can exist in two forms with opposite curvature, one with flagella pointing towards the center of curvature [“*flag-in*”] with a relatively large spacing between cells, and another with the opposite arrangement [“*flag-out*”] with more tightly-packed cells. Transformations from *flag-in* to *flag-out* can be triggered by darkness, and occur in ∼ 10 s. Compared to the *flag-out* form, the *flag-in* state has limited motility and is better suited to filter-feeding. It was conjectured [8] that the darkness-induced transition to the more motile form is a type of photokinesis.

**FIG. 1.**
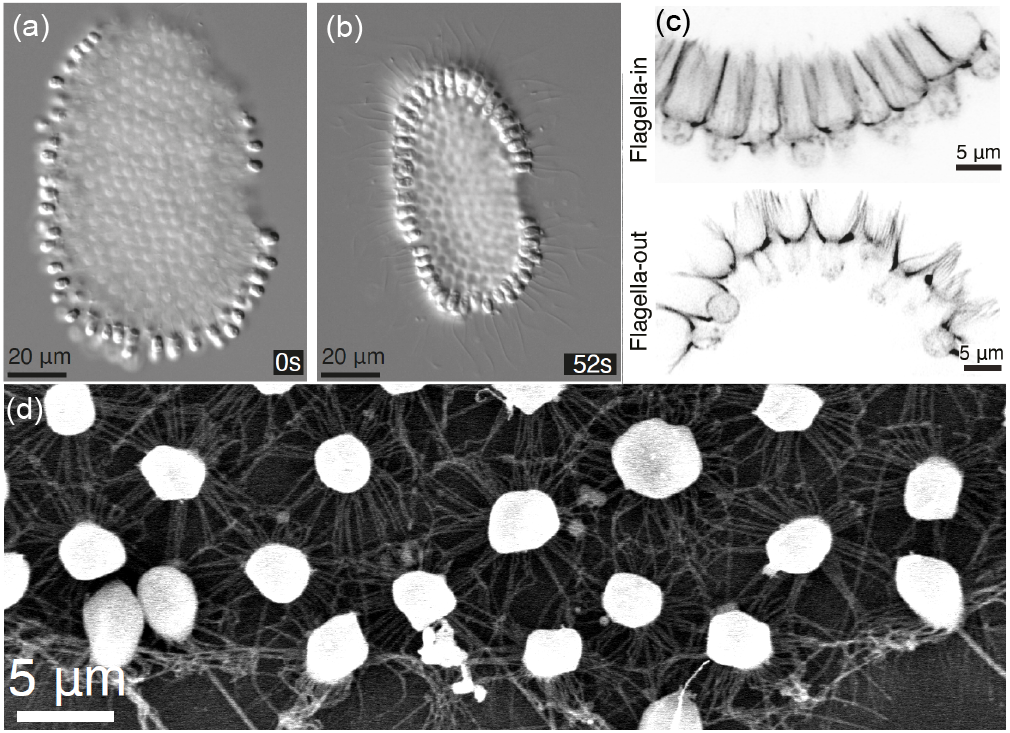
The multicellular choanoflagellate *Choanoeca flexa*. Top views of (a) *flag-in* and (b) *flage-out* states at times relative to removal of light. (c) Close-up of the collar connections in the two states. (d) Electron micrograph showing round white cell bodies connected by microvilli. Adapted from [8].

As a first step toward understanding principles that govern the behavior of such a novel organism as *C. flexa*, we analyze two models for these shape-shifting structures. First, the fluid mechanics are studied by representating the cell raft as a collection of spheres distributed on a hemispherical surface, with nearby point forces to represent the action of flagella. Such a model has been used to describe the motility of small sheet-like multicellular assembles such as the alga *Gonium* [10]. The motility and filtering flow through these rafts as a function of cell spacing and curvature explain the observed properties of *C. flexa*. Second, abstracting the complex elastic interactions between cells to the simplest connectivity, we show that a model based on linear elasticity at the microscale produces bistability on the colony scale.

## Fluid mechanics of feeding and swimming

The cells in a *C. flexa* raft are ellipsoidal, with major and minor axes *a* ∼ 4 *µ*m and *b* ∼ 3 *µ*m, with a single flagellum of length 2*L* ∼ 24.5 *µ*m and radius *r* ∼ 0.5 *µ*m beating with amplitude *d* ∼ 2.3 *µ*m and frequency *f* ∼ 44 Hz [11], sending bending waves away from the body. A cell swims with flagellum and collar rearward; the body and flagellum comprise a “pusher” force dipole. From resistive force theory [13] we estimate the flagellar propulsive force to be *F* ∼ 2*L* (*ζ*_⊥_ − *ζ*_∥_) (1−*β*)*fλ* ∼ 8 pN, where *β* is a function of the wave geometry, *λ* ∼ 12 *µ*m is the wavelengths along the direction of the bending wave [11], *ζ*_⊥_ and *ζ*_∥_ are transverse and longitudinal drag coefficients, *ζ*_⊥_ ∼ 2*ζ*_∥_ ∼ 4*πµ/* ln(2*L/r*), with *µ* the fluid viscosity. These features motivate a computational model in which *N* identical cells in a raft have a spherical body of radius *a* and a point force 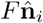 acting on the fluid a distance *L* from the sphere center, oriented along the vector 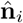 that represents the collar axis [Fig. 2(a,b)]. An idealization of a curved raft involves placing those spheres on a connected subset of the vertices of a *geodesic* icosahedron (one whose vertices lie on a spherical surface) of radius *ρ* ≫ *a*; the area fraction Φ of the sheet occupied by cells scales as Φ ∼ *N*(*a/ρ*)^2^. The pentagonal neighborhoods within the geodesic icosahedron serve as topological defects that allow for smooth large-scale surface curvature [14]. Importantly, confocal imaging of *C. flexa* colonies shows that a significant fraction (∼ 0.25) of the cellular neighborhoods defined by the microvilli connections are pentagonal [11], and earlier work on *C. perplexa* [9] also found non-hexagonal packing. We use the geodesic icosahedron {3, 5+}_(3,0)_ in standard notation [15], with 92 total vertices, and take patches with *N* = 58 for computational tractability. The vectors 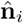 point towards (away from) the icosahedron center in the *flag-in* (*flag-out*) forms (Fig. 2(a)). A deformation of the sheet to a new radius *ρ*′, at fixed Φ, requires the new polar angle 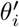 of a cell with respect to the central axis of the sheet be related to its original angle *θ*_*i*_ via 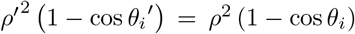. We define the scaled force offset length *ℓ* = *L/a* ∼ 3 and sheet radius *R* = *ρ*′/*a* ≳ 6, and take *R >* 0 in the *flag-in* state.

**FIG. 2.**
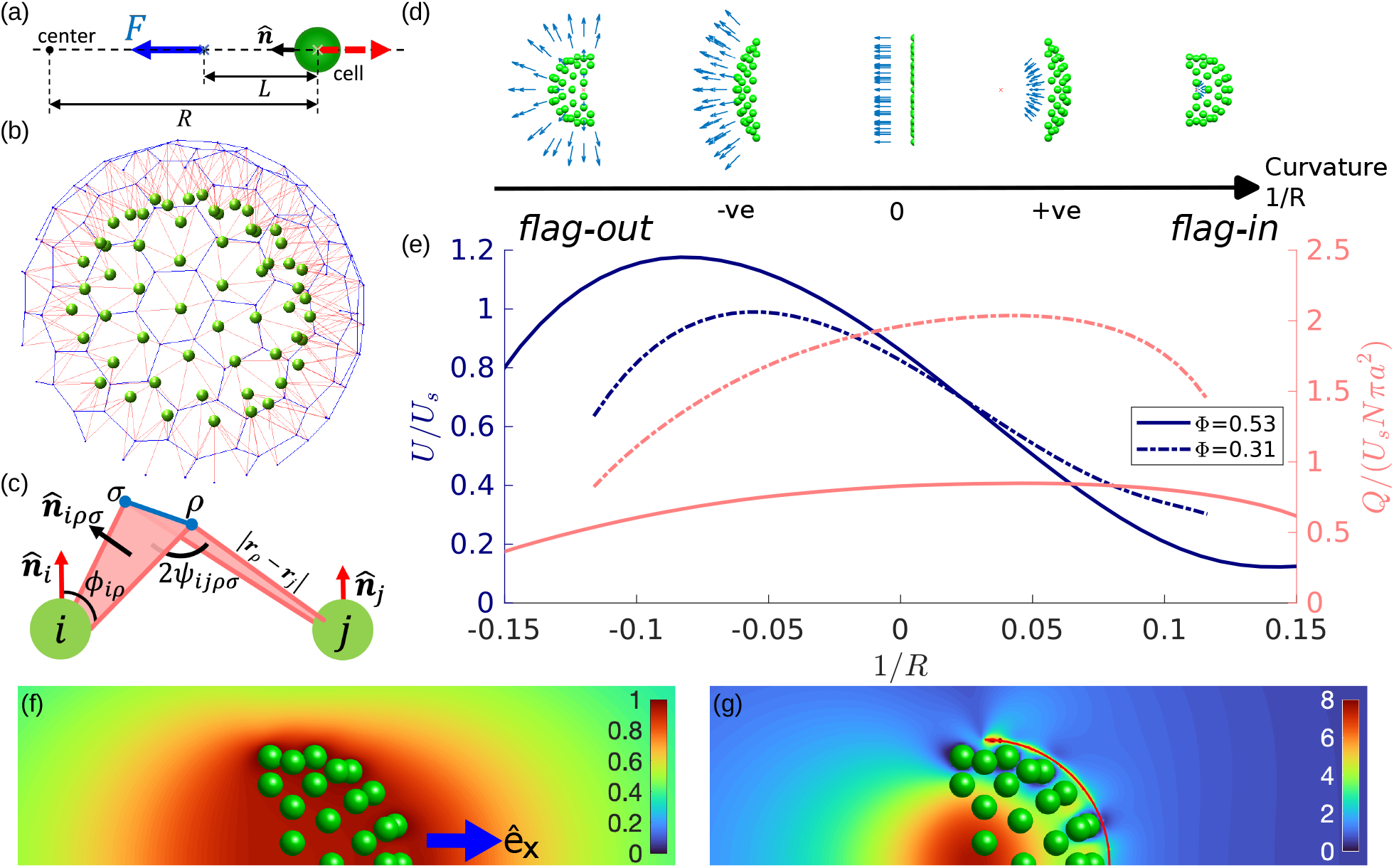
Models for *C. flexa*. (a) Cell body and flagellar force in the *flag-in* state. (b,c) Mechanical model of interconnecting microvilli in rafts; cells (green spheres, not to scale) are at the vertices of a geodesic icosahedron. Blue arrows indicate flagella forces, red segments represent microvilli, blue dots the microvilli tips, and blue lines the collar-collar interface. (b) Connectivity of the whole raft. (c) Connections between two cells. Effect of curvature 1/*R* on (d) geometry of raft and (e) swimming speed *U* (blue) and flow rate *Q* (red) passing through *S*_*f*_ at constant. (f,g) Cross section of the disturbance flows **ũ**_*d*_ and **u**_*f*_ around the a raft (Φ = 0.31, *R* = 8.61) in the reciprocal problems for calculating *U* and *Q*.

Images of many colonies of *C. flexa* [11] show that the packing fraction in the *flag-out* state Φ_out_ = 0.47 ± 0.06, considerably less than both the maximum packing fraction 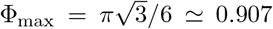 for a hexagonal array of spheres in a plane, and the estimated maximum packing fraction 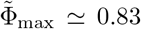 for circles on a sphere [16]. The packing fraction in the *flag-in* state is Φ_in_ = 0.34 ± 0.03, and we use the extremes Φ_out_ = 0.53 and Φ_in_ = 0.31 as representative values to explore the consequences of the differences between the two forms.

Consider first an isolated force-free spherical cell at the origin moving at velocity *U*_*s*_**ê**_*x*_ with point force − **F** = − *F***ê**_*x*_ at − *L***ê**_*x*_ acting on the fluid and its reaction force **F** acting on the cell. The cell experiences Stokes drag −*ζ*_*s*_*U*_*s*_**ê**_*x*_, where *ζ*_*s*_ = 6*πµa*, and a disturbance drag − *D***ê**_*x*_ arising from the disturbance flow created by the point force. By the reciprocal theorem [17], the disturbance drag is *D* = **F** · **ũ**_*d*_(−*L***ê**_*x*_)/*Û*, where **ũ**_*d*_(**r**) is the disturbance flow created when the cell is dragged along **ê**_*x*_ with unit speed *Û*. Force balance then yields the single-cell swimming speed *U*_*s*_ ≡ (*F/ζ*_*s*_)[1 − 3/(2*ℓ*) + 1/(2*ℓ*^3^)]. Thus, the closer the point force is to the cell (i.e., the smaller is *ℓ*), the more drag the cell experiences and the slower is *U*_*s*_. Setting *ℓ* = 3 yields *U*_*s*_ ∼ 118 *µ*m/s, consistent with observations.

This intuitive picture extends to a raft of cells. As the raft moves at velocity *U* **ê**_*x*_, it experiences a Stokes drag −*ζU* **ê**_*x*_. The disturbance flow created by the point forces 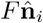 acting at 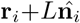, produces a disturbance drag 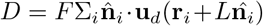, where **u**_*d*_ is the (dimensionless) disturbance flow from the raft when it is dragged along **ê**_*x*_ with unit speed. Force balance then yields

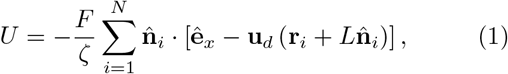

where 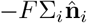 is the sum of reaction forces propelling the raft along **ê**_*x*_, and where **u**_*d*_ has been rendered dimensionless by the unit speed. In practice, we compute **u**_*d*_ and *ζ*using a Boundary Element Method [10]. Because of the curved geometry, point forces are closer to neighboring cells in the *flag-in* state than in the *flag-out* state. Thus, as in Fig. 2(d), for a geometry with a given |*R*|, the *flag-in* state has a larger disturbance drag than the *flag-out* state, and a smaller speed *U*.

The difference in swimming speed between the two states can also be explained in terms of 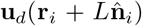 in (1). Figure 2(f) shows that **u**_*d*_ inside the raft is close to **ê**_*x*_ because of the curved geometry and screening effects. Hence, *U* is small when the point forces are inside. Meanwhile, **u**_*d*_ outside decays with the distance from the raft, so *U* is large when 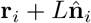 is outside.

Previous work on filter-feeding in choanoflagellates focused first on the far-field limit based on a stresslet description [18], but later work showed near-field effects can significantly affect capture rates [19]. To estimate the filter-feeding flux *Q* passing through a colony of *C. flexa*, we measure, in the body frame, the flux passing through the surface *S*_*f*_ projected a distance of 1.2*a* from the cell center along 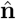, as in Fig. 2(g). By the reciprocal theorem, *Q* can be written in terms of the disturbance flow **u**_*f*_ around a stationary raft and the hydrodynamic forces **F**_*f*_ on the raft when the surface *S*_*f*_ applies a unit normal pressure 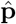 on the fluid,

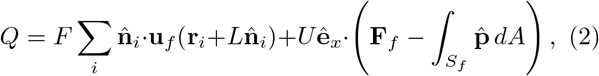

where **u**_*f*_ and **F**_*f*_ acquire the units of velocity/pressure and area, respectively, by scaling with 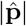. Numerical results in Fig. 2(d) show that the flux due to point forces 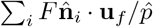 strongly dominates *Q*. Therefore, the difference in *Q* between the two states can be explained by 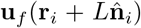 (Fig. 2(g)). To maintain incompressibility under pressure 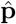, the disturbance flow **u**_*f*_ is much stronger inside the raft than outside. Hence, point forces placed inside the raft pump more flow through the raft than when placed outside.

Figure 2(d) shows the effect of changes in the raft curvature and packing fraction. There is one *R* that maximizes swimming speed in the *flag-in* state and another one that maximizes feeding flux in the *flag-out* state. This arises from a balance between the screening effect mentioned above and the alignment of forcing. In the *flag-out* state, an initial decrease in curvature aligns the forcing direction with the swimming direction, increasing swimming speed, but a further reduction in curvature reduces the screening effect as cells are now more spread out in the plane orthogonal to the swimming direction. A similar argument applies to the flow rate maximum in the *flag-in* state. Comparing these maxima, Fig. 2(d) shows that a spread-out colony results in more flux, while a closely-packed colony results in faster motility. Thus, through the interconversion between the two states, *C. flexa* takes advantage of the hydrodynamics effect of the curved geometry for efficient filter-feeding and swimming.

## Mechanics of inversion

Detailed studies suggest that inversion requires an active process within each cell, likely driven by contraction of an F-actin ring at the apical pole through the action of myosin [8]. Thus, a full treatment would address the complex problem of elastic filaments responding to the apical actomyosin system and adhering to each other. We simplify this description by considering as in Fig. 2(e) that each cell *i*, located at **r**_*i*_ and surrounded by *m*_*i*_ neighbors, has *m*_*i*_ rigid, straight filaments emanating from it. Two filaments from neighboring cells *i* and *j* meet at vertex *ρ* located at **r**_*ρ*_, with *ϕ*_*iρ*_ the angle between **r**_*ρ*_ − **r**_*i*_ and the cell normal vector 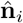. Any two adjacent filaments emanating from cell *i*, and which meet neighboring filaments at vertices *ρ* and *σ*, define a plane whose normal 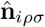 points toward the apicobasal axis 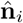. That normal and its counterpart 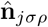 on cell *j* determine the angle 2*ψ*_*ijρσ*_ between the two planes.

As above, we use the geodesic icosahedron to define the cell positions and thus determine the filament net-work connecting neighboring cells. The two sets of angles {*ϕ*} and {*ψ*} are used to define a Hookean elastic energy that mimics the elasticity of the microvilli, allowing for preferred intrinsic angles *ϕ*_0_ and *ψ*_0_ that encode the effects of the apical actomyosin system on the microvilli and the geometry of microvilli adhesion. Allowing also for stretching away from a rest length *ℓ*_0_, the energy is

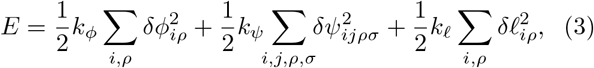

where *δϕ*_*iρ*_ = *ϕ*_*iρ*_ − *ϕ*_0_, *δψ*_*ijρσ*_ = *ψ*_*ijρσ*_ − *ψ*_0_, and *ℓ*_*iρ*_ = |**r**_*i*_ − **r**_*ρ*_| − *ℓ*_0_. The moduli *k*_*ϕ*_, *k*_*ψ*_ and *k*_*ℓ*_ and quantities *ϕ*_0_, *ψ*_0_ and *ℓ*_0_ are assumed constant for all cells.

The energy (3) is intimately tied to the lattice geometry of the raft. If the cells are arranged in a hexagonal lattice (*m*_*i*_ = 6) the system of filaments can achieve *E* = 0 by setting all cell-collar angles to *ϕ*_0_, all collar-collar interface angles to *ψ*_0_, and *ϕ*_0_ = *ψ*_0_. This corresponds to a flat sheet. Increasing *ϕ*_0_ = *ψ*_0_ leads to uniform, isotropic sheet expansion. In a non-planar raft, curvature is introduced through topological defects (*m*_*i*_ ≠ 6), such as pentagons, and mismatch between the local values of *ψ* and *ϕ*. While pentagonal defects are known to cause out-of-plane buckling in crystal lattices [14], they do not by themselves select a particular *sign* of the induced curvature. Thus, there is inherent bistability in the cellular raft that can be biased by changes in the geometry of the out-of-plane filaments, somewhat akin to the role of “apical constriction” in the shapes of epithelia [20].

For the case of two cells lying in a plane, each with two filaments, and with one vertex between them, if *ϕ* = *ϕ*_0_, *ψ* = *ψ*_0_, and *r* = *ℓ*_0_, then the filament tips lie on a circle of radius *R*_0_ = 1/*C*_0_, where *C*_0_ = sin(*ψ*_0_ − *ϕ*_0_)/*ℓ*_0_ sin *ϕ*_0_. While, in general, the equilibrium state of a curved raft will not have *ϕ*_*iρ*_ = *ϕ*_0_, *ψ*_*ijρσ*_ = *ψ*_0_ and *r*_*iρ*_ = *ℓ*_0_ everywhere, we may nevertheless use this relationship to define a proxy for the average curvature of the raft. Recognizing that in numerical studies stretching effects are small, we ignore variations in *r*_*iρ*_ and define *C* = sin(⟨*ψ*⟩ − ⟨*ϕ*⟩)/*ℓ*_0_ sin(⟨*ϕ*⟩), where ⟨·⟩ is an average over cells and vertices. The colony is in the *flag-in* (*flag-out*) state when *C >* 0 (*C <* 0).

The simplest model of raft dynamics localizes the viscous drag to the individual cell and vertex positions **r**_*γ*_ according to a gradient flow *ζ*∂_*t*_**r**_*γ*_ = − ∂*E/*∂**r**_*γ*_ driven by the force derived from (3). We solve this dynamics numerically with forward integration. Since the 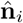 are constrained to have unit length, they are normalized after each step in the direction of the negative gradient, making the dynamical algorithm follow a projected gradient descent [21]. Via a rescaling of time we may set one of the elastic constants to unity (say, *k*_*ϕ*_) and need only consider the ratios *K*_*ψ*_ = *k*_*ψ*_/*k*_*ϕ*_ and *K*_*ℓ*_ = *k*_*ℓ*_/*k*_*ϕ*_.

Interconversion between the *flag-in* and *flag-out* states is shown in Figs. 3(a-d) following an abrupt change in the preferred angle pair (*ϕ*_0_, *ψ*_0_) that crossing the line of equality *ψ*_0_ = *ϕ*_0_ that divides the states (Fig. 3(e)), as during raction/relaxation of the F-actin ring in response to a stimulus. The intermediate shapes exhibit a ring of inflection points similar to those seen in experiments on *C. flexa* and also in the inversion the algae *Pleodorina* [3] and larger species [22, 23]. Tracking the energy as each of the two equilibria is achieved, the picture that emerges in Fig. 3(f) is evolution on a double-well potential energy landscape as a biasing field is switched in sign.

**FIG. 3.**
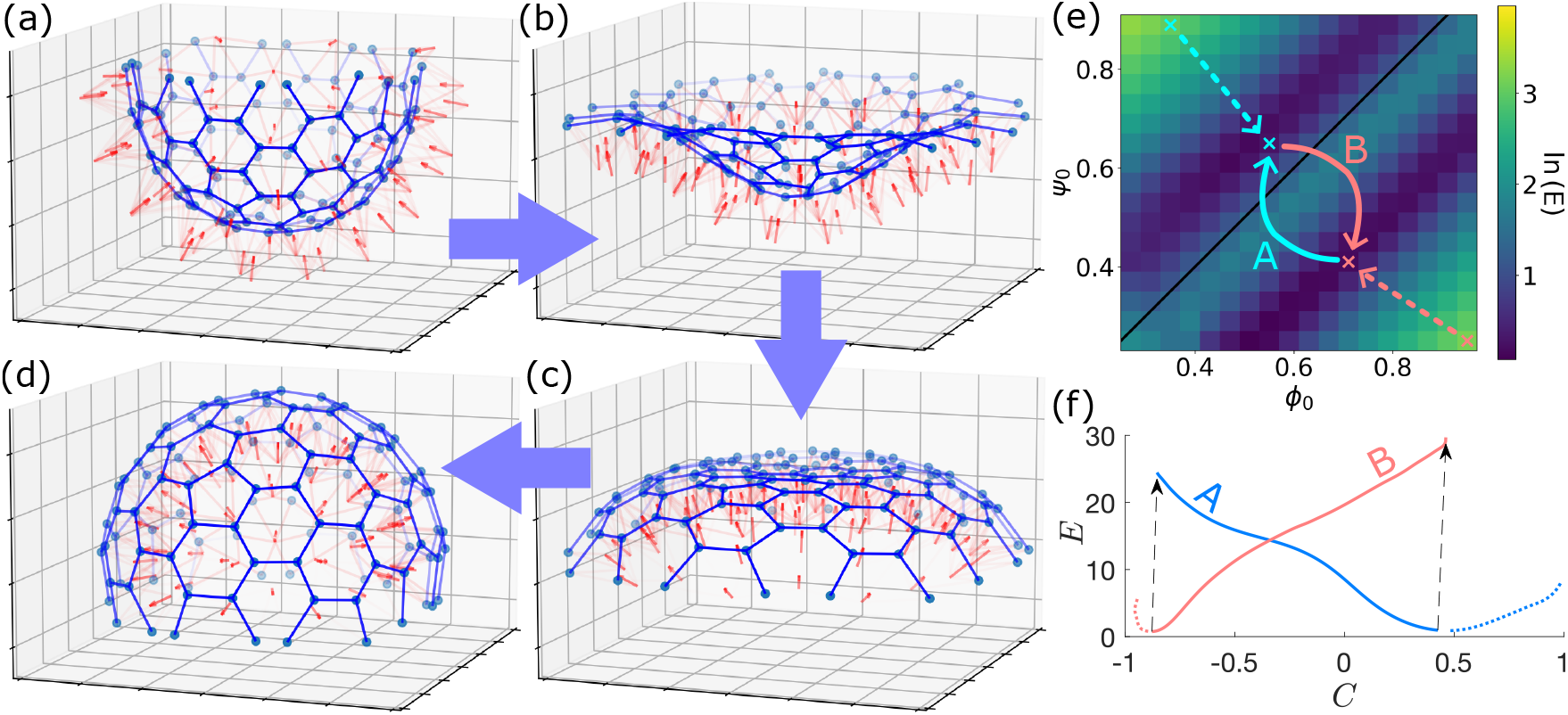
Inversion dynamics from numerical studies. (a)-(d) A colony, initially at a hemispherical minimum with (*ϕ*_0_, *ψ*_0_) = (0.55, 0.65), inverts after a change to (0.71, 0.41), with *ℓ*_0_ = 0.5, *K*_*ψ*_ = 2 and *K*_*ℓ*_ = 5. Connections between collar vertices are shown in blue, apicobasal axes as red arrows at cell body positions. (e) Minimum energy *E* in the *ψ*_0_ *− ϕ*_0_ plane, where *C <* 0 (*flag-in*) above the black line *ψ*_0_ = *ϕ*_0_, and *C >* 0 (*flag-out*) below. (*f*) Evolution of *E* vs *C* as the colony relax towards a minimum energy state after instantaneous changes in (*ϕ*_0_, *ψ*_0_) shown by the dotted and solid red and blue lines in (*e*).

We have shown that simple models can explain the swimming, feeding, and inversion of the recently discovered multicellular choanoflagellate *C. flexa* [8]. These results suggest further exploration on a possible continuum description of the sheets, fluid-structure interactions during locomotion, dynamics of photokinesis, and developmental processes of these remarkable organisms.

We gratefully acknowledge Gabriela Canales and Tanner Fadero for high speed imaging assistance and Kyriacos Leptos for comments and suggestions. This work was supported in part by a Research Fellowship from Peterhouse, Cambridge (LF), a Churchill Scholarship (AK), JSPS Kakenhi (TI), The John Templeton Foundation and Wellcome Trust Investigator Grant 207510/Z/17/Z (REG). For the purpose of open access, the authors have applied a CC BY public copyright license to any Author Accepted Manuscript version arising from this submission.

## SUPPLEMENTAL MATERIAL

This file contains additional experimental results on flagellar dynamics and geometry of *C. flexa*.

### I. VIDEO IMAGING AND ANALYSIS

#### A. Supplementary Videos

*C. flexa* sheets were imaged in FluoroDishes (World Precision Instruments FD35-100) by differential interference contrast (DIC) microscopy using a 40× (water immersion, C-Apochromat, 1.1 NA) Zeiss objective mounted on a Zeiss Observer Z.1 with a pco.dimax cs1 camera.

Flagellar characteristics reported in Table I were obtained as follows. Beat frequencies were determined by averaging over five cycles for each of twenty randomly selected cells. All other measurements are averages over ten randomly selected cells. The comparatively large wavelength for *flag-out* sheets may be due in part to the fact that the flagellar waveform in that state is not sinusoidal, and its wavelength is thus less well defined than in the *flag-in* state.

**FIG. S1.**
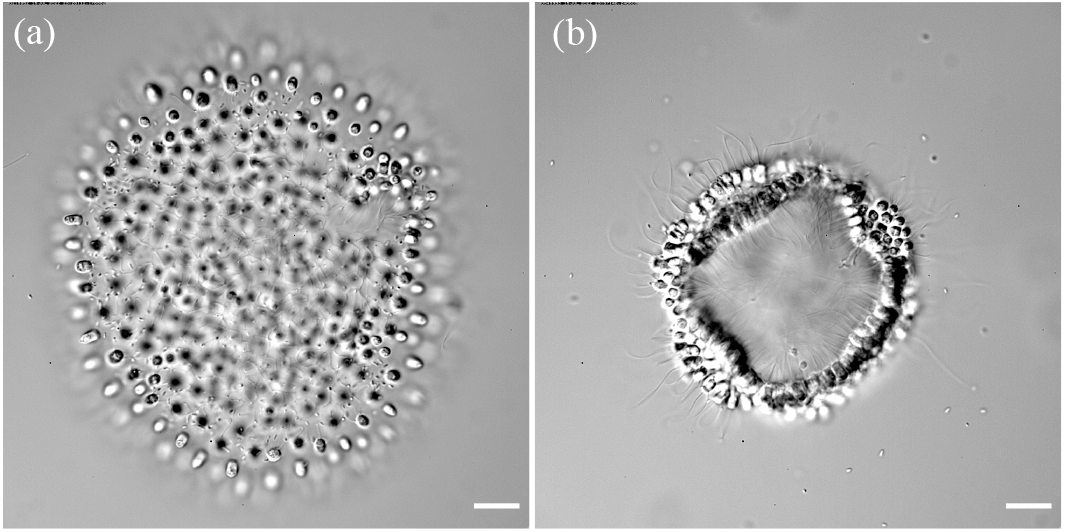
Flagellar dynamics in *C. flexa* colonies. (a) Snapshot from Video 1, a high speed recording of a *flag-in* sheet used to determine flagellar characteristics reported in Table I. Movie is set to play at 17× slower than real time. (b) As in (a), but for the *flag-out* state in Video 2. Scale bars are 10 *µ*m.

**TABLE I.**
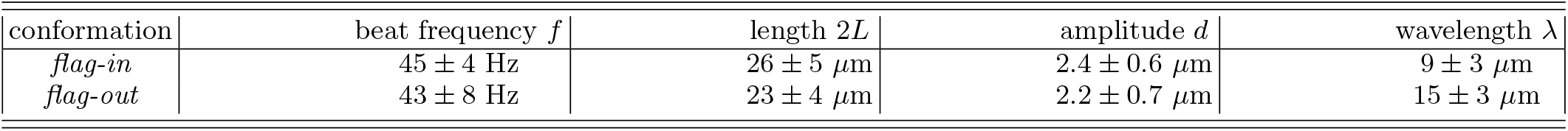
Measurements of flagellar characteristics for the *flag-in* and *flag-out* sheets in Videos S1 and S2. Uncertainties reported are standard deviations.

| conformation | beat frequency $f$ | length $2L$ | amplitude $d$ | wavelength $\lambda$ |
| --- | --- | --- | --- | --- |
| *flag-in* | $45 \pm 4\ \text{Hz}$ | $26 \pm 5\ \mu\text{m}$ | $2.4 \pm 0.6\ \mu\text{m}$ | $9 \pm 3\ \mu\text{m}$ |
| *flag-out* | $43 \pm 8\ \text{Hz}$ | $23 \pm 4\ \mu\text{m}$ | $2.2 \pm 0.7\ \mu\text{m}$ | $15 \pm 3\ \mu\text{m}$ |

#### B. Estimating the propulsive force from the flagella

In the fluid mechanics model of the *C. Flexa* raft, we approximate the flagella beating as an effective propulsive point force **F** acting in the direction 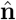. The magnitude of this force *F* can be approximated using the resistive force theory [S1] as

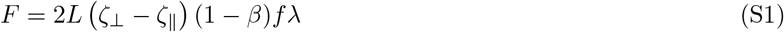

where 2*L* is the flagella length, *f* the beat frequency and *λ* the projected wavelength in the direction of the traveling sinusoidal wave (i.e.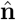), the values of which are listed in Table I. Meanwhile,

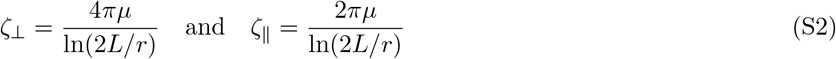

are the transverse and longitudinal drag coefficients of a cylindrical filament of radius *r*, approximated using the resistive force theory, and *β* is a coefficient that depends on the flagella waveform. Although the flagella waveform is not necessarily sinusoidal, in the absence of better measurements, the value of *β* is approximated, assuming the flagella takes a sinusoidal waveform *f* (*x*) with wavelength *λ* and amplitude *d* (Table I), as

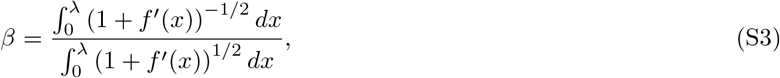

which can be found by numerically. In the limiting of 2*πd/λ* ≪ 1, *β* ≈ 2*π*^2^(*d/λ*)^2^.

### II. CONFOCAL IMAGING

Sheets in Figs. S2 and S3 were fixed and stained with FM1-43FX or with Alexa 488-phalloidin as in [S2]. Sheets were imaged on Zeiss LSM 880 with AiryScan using a 63x, 1.4 NA C Apo oil immersion objective (Zeiss). Z-projections were generated with Fiji [S3]. Packing fraction was estimated by projecting cell bodies located within the same plane in a locally flat portion of the sheet and by manually outlining the border of the colonies (red dotted line in Fig. S2).

**FIG. S2.**
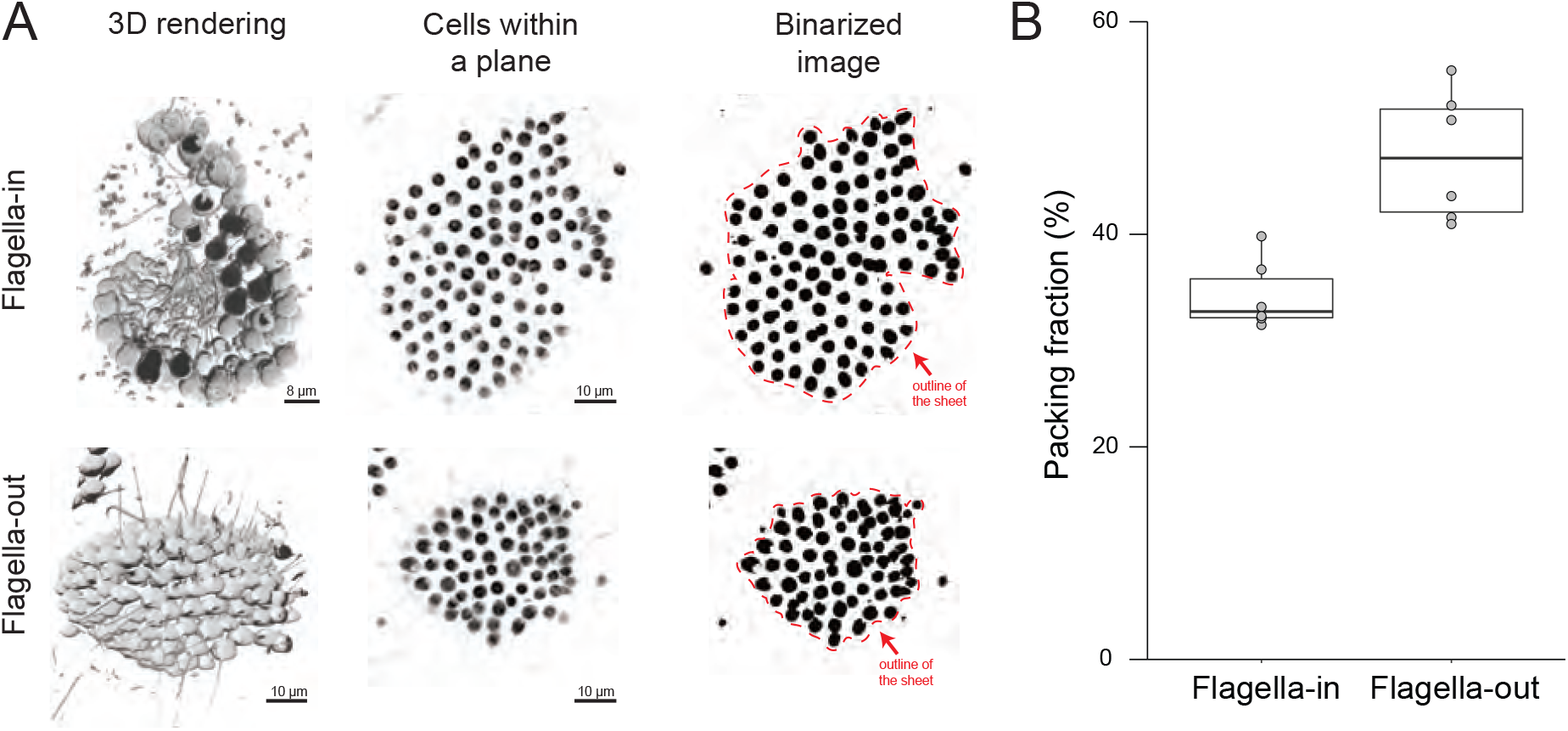
Packing fraction in flagella-in and flagella-out *C. flexa* colonies. (a) 3D stacks (left column) of sheets stained with the fluorescent membrane marker FM 1-43 FX were imaged by confocal microscopy to determine sheet morphology. Packing fraction was computed by doing a Z-projection of a locally flat portions of individual sheets (middle column) and generating a binarized image in which the area occupied by individual cells appears black (right column). Packing fraction is the ratio of the area occupied by cells to the total colony area within that plane (area within the red dotted line). (b) Boxplot depicting packing fraction values for 6 sheets with flagella out and 6 sheets with flagella-in. p=0.2% by the Mann-Whitney test.

Packing fraction was then computed as the ratio of the area occupied by cells (black area in binarized image in Fig. S2) to the total area occupied by the colony (area within the red dotted line in Fig. S2). Polygonal collar borders in Fig. S3 were manually outlined and colored with Adobe Illustrator 27.3.1 (2023). Hexagonal and pentagonal outlines were counted in 9 colonies and counts are reported in Table II.

**TABLE II.** Numbers of hexagons and pentagons in representative sections of 9 colonies of *C. flexa*.

| colony # | 1 | 2 | 3 | 4 | 5 | 6 | 7 | 8 | 9 | total |
| --- | --- | --- | --- | --- | --- | --- | --- | --- | --- | --- |
| hexagons | 4 | 4 | 2 | 8 | 8 | 15 | 6 | 6 | 8 | 61 |
| pentagons | 4 | 0 | 1 | 2 | 2 | 7 | 2 | 3 | 2 | 23 |

**FIG. S3.**
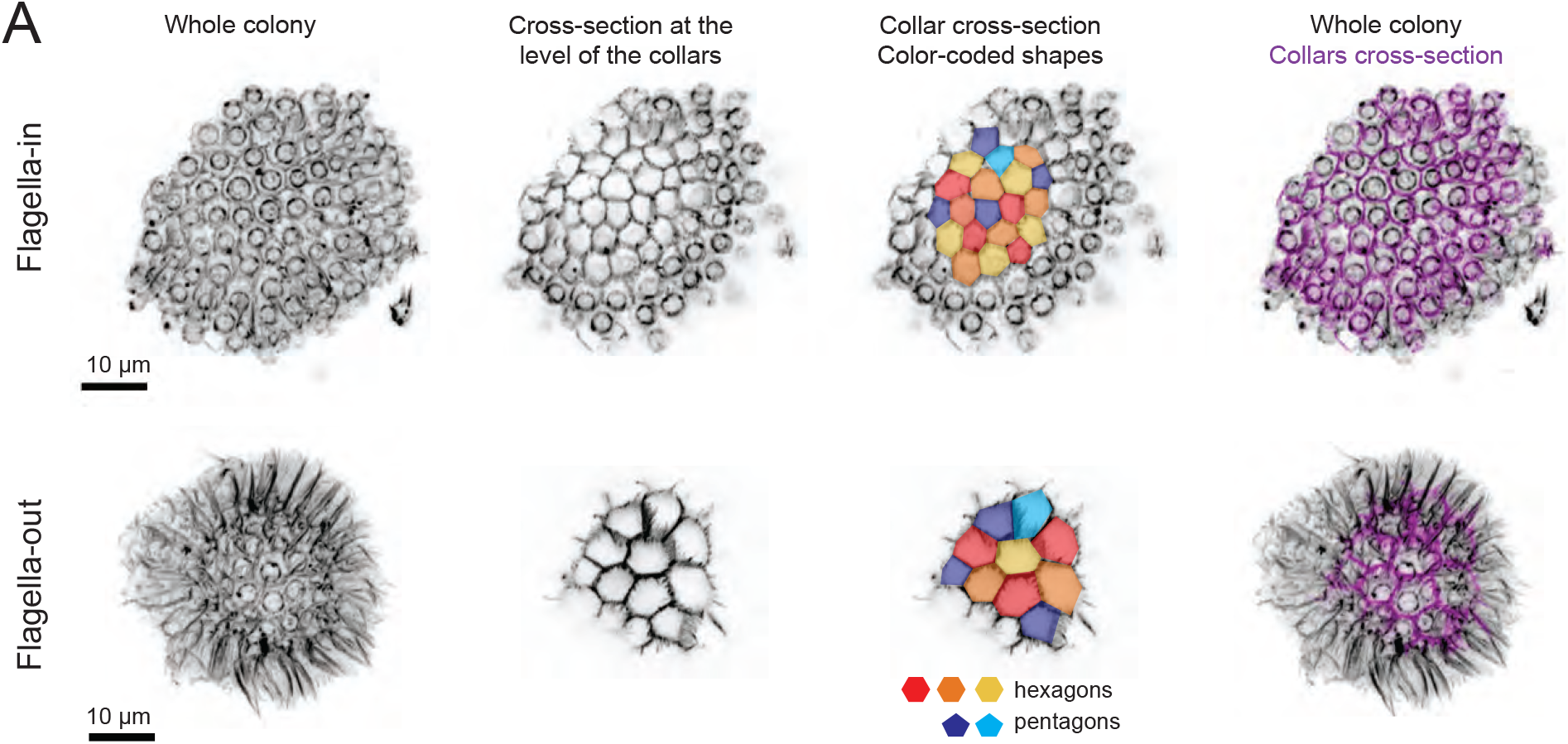
Hexagonal and polygonal collar outlines within flagella-in and flagella-out *C. flexa* colonies. F-actin within colonies was stained with fluorescent phalloidin, which outlines the microvillous collars linking cells, as well as the actin cytoskeleton within the cell body. Left column: whole-colony Z-projections. Middle left: Z-projections of a few planes intersecting many collars, showing the polygonal outlines of collar contacts. Middle right: color-coded polygons showing a majority of hexagons and a minority of pentagons. Right: whole-colony Z-projection with the plane containing most collars outlined in purple.

## Notes

### Competing Interest Statement

The authors have declared no competing interest.

## References

[1] L. Solnica-Krezel and D.S. Sepich, Gastrulation: Making and Shaping Germ Layers, Annu Rev. Cell Dev. Biology 28, 687–717 (2012).

[2] A. Goriely, D.E. Moulton and R. Vandiver, Elastic cavitation, tube hollowing, and differential growth in plants and biological tissues, EPL 91, 18001 (2010).

[3] S. Höhn and A. Hallmann, Distinct shape-shifting regimes of bowl-shaped cell sheets–embryonic inversion in the multicellular green alga Pleodorina, BMC Dev. Biol. 16, 35 (2016).

[4] S. Höhn, A.R. Honerkamp-Smith, P.A. Haas, P. Khuc Trong, and R.E. Goldstein, Dynamics of a Volvox embryo turning itself inside out, Phys. Rev. Lett. 114, 178101 (2015).

[5] T. Brunet and N. King, The Origin of Animal Multicellularity and Cell Differentiation, Dev. Cell 43, 124–140 (2017).

[6] S.R. Fairclough, M.J. Dayel, and N. King, Multicellular development in a choanoflagellate, Curr. Biology 20, R875–R876 (2010).

[7] B.T. Larson, T. Ruiz-Herrero, S. Lee, S. Kumar, L. Mahadevan and N. King, Biophysical principles of choanoflagellate self-organization, Proc. Natl. Acad. Sci. USA 117, 1303–1311 (2020).

[8] T. Brunet, B.T. Larson, T.A. Linden, M.J.A. Vermeij, K. McDonald, and N. King, Light-regulated collective contractility in a multicellular choanoflagellate, Science 366, 326–33 (2019).

[9] As noted previously [8], the behavior of C. flexa is similar to the species C. perplexa studied earlier: B.S.C. Leadbeater, Life-history and ultrastructure of a new marine species of Proterospongia (Choanoflagellida), J. Mar. Biol. Ass. U.K. 63, 135–160 (1983).

[10] H. de Maleprade, F. Moisy, T. Ishikawa, and R.E. Goldstein, Motility and phototaxis in Gonium, the simplest differentiated colonial alga Phys. Rev. E 101, 022416 (2020).

[11] See Supplemental Material at http://link.aps.org/supplemental/xxx for further experimental results, which includes Ref. [12].

[12] J. Schindelin, I. Arganda-Carreras, E. Frise et al. Fiji: an open-source platform for biological-image analysis. Nat Methods 9, 676–682 (2012).

[13] E. Lauga, The Fluid Dynamics of Cell Motility (Cambridge University Press, Cambridge, UK, 2020).

[14] H.S. Seung and D.R. Nelson, Defects in flexible membranes with crystalline order, Phys. Rev. A 38, 1005–1018 (1988).

[15] M. Wenninger, Spherical Models (Cambridge University Press, Cambridge, UK, 1979).

[16] B.W. Clare and D.L. Kepert, The Optimal Packing of Circles on a Sphere, J. Math. Chem. 6, 325–349 (1991).

[17] S. Kim and S. J. Karrila, Microhydrodynamics: Principles and Selected Applications (Butterworth-Heinemann, Boston, US, 1991)

[18] Roper, M. and Dayel, M.J. and Pepper, R.E. and Koehl, M.A.R., Cooperatively Generated Stresslet Flows Supply Fresh Fluid to Multicellular Choanoflagellate Colonies, Phys. Rev. Lett. 110, 228104 (2013).

[19] J.B. Kirkegaard and R.E. Goldstein, Filter-feeding, nearfield flows, and the morphologies of colonial choanoflagellates, Phys. Rev. E 94, 052401 (2016).

[20] E. Hannezo, J. Prost, J.-F. Joanny, Theory of epithelial sheet morphology in three dimensions, Proc. Natl. Acad. Sci. USA 111, 27–32 (2014).

[21] B. Eicke, Iteration methods for convexly constrained illposed problems in Hilbert space Num. Func. Anal. Opt. 13, 413–429 (1992).

[22] G.I. Viamontes and D.L. Kirk, Cell shape changes and the mechanism of inversion in Volvox, J. Cell Biol. 75, 719–730 (1977).

[23] P.A. Haas, S. Höhn, A.R. Honerkamp-Smith, J.B. Kirkegaard, and R.E. Goldstein, The Noisy Basis of Morphogenesis: Mechanisms and Mechanics of Cell Sheet Folding Inferred from Developmental Variability, PLOS Biol. 16, e2005536 (2018).

## References

[S1] E. Lauga, The Fluid Dynamics of Cell Motility (Cambridge University Press, Cambridge, UK, 2020), §7.1.3-7.1.4.

[S2] T. Brunet, B.T. Larson, T.A. Linden, M.J.A. Vermeij, K. McDonald, and N. King, Light-regulated collective contractility in a multicellular choanoflagellate, Science 366, 326–33 (2019).

[S3] J. Schindelin, I. Arganda-Carreras, E. Frise et al. Fiji: an open-source platform for biological-image analysis. Nat Methods 9, 676–682 (2012).

